# Development of a potent and protective germline-like antibody lineage against Zika virus in a convalescent human

**DOI:** 10.1101/661918

**Authors:** Fei Gao, Xiaohe Lin, Linling He, Ruoke Wang, Han Wang, Xuanling Shi, Fuchun Zhang, Chibiao Yin, Linqi Zhang, Jiang Zhu, Lei Yu

**Affiliations:** Comprehensive AIDS Research Center, Collaborative Innovation Center for Diagnosis and Treatment of Infectious Diseases, Department of Basic Medical Sciences, School of Medicine, Tsinghua University, Beijing, China; Department of Integrative Structural and Computational Biology, Department of Immunology and Microbiology, The Scripps Research Institute, La Jolla, CA, United States; Guangzhou Eighth People’s Hospital, Guangzhou Medical University, Guangzhou, China

**Keywords:** Zika virus infection, Guillain–Barré syndrome, microcephaly, neutralizing antibody, antibody repertoire, next-generation sequencing

## Abstract

Zika virus (ZIKV) specific neutralizing antibodies hold a great promise for antibody-based interventions and vaccine design against ZIKV infection. However, their development in infected patients remain unknown. Here, we report on the dynamic development of a potent and protective ZIKV-specific human antibody ZK2B10 initially isolated from a ZIKV convalescent individual using next-generation sequencing (NGS). The unbiased repertoire analysis showed dramatic changes in many families of heavy and light chain variable regions. However, lineage tracing of ZK2B10 revealed limited somatic hypermutation throughout the 12 months since the onset of symptom. In particular, NGS-derived germline-like somatic variants neutralized and protected mice from lethal challenge of ZIKV without detectable cross-reactivity with Dengue virus (DENV). Site-directed mutagenesis identified two residues within λ chain, N31 and S91 that are essential to the functional maturation. The dynamic features unveiled here will assist us to better understand the pathogenesis of ZIKV infection and inform rational design of vaccines.

**Author summary:** Recently emerged ZIKV is associated with severe neurological complications such as Guillain–Barré syndrome in adults and congenital microcephaly in newborns. No ZIKV-specific therapeutics or vaccines are currently available. We and others have identified a number of neutralizing antibodies capable of protecting experimental animals from ZIKV infection. However, the development of these potent antibodies during ZIKV natural infection remains unknown. Here, we report on the longitudinal analysis of one such antibody ZK2B10 using next-generation sequencing (NGS), bioinformatics and functional analysis. We found that the ZK2B10 germline-like antibodies possess strong neutralizing activity *in vitro* and impressive protectivity against lethal ZIKV infection *in vivo*. These findings suggest that the potent and protective antibody response against ZIKV can be generated within relative short term with high germline identity which provide great hope and promise for successful vaccine development against ZIKV.

## Introduction

Zika virus (ZIKV), a member of the *Flavivirus genus* of the *Flaviviridae* family, is an emerging mosquito-borne pathogen. ZIKV is closely related to other flavivirus such as dengue (DENV 1, 2, 3 and 4), yellow fever (YFV), West Nile (WNV), Japanese encephalitis (JEV), and tick-borne encephalitis (TBEV) viruses [1]. Since ZIKV was first identified in 1947 among rhesus macaques of Uganda Zika forest, its new variants have become increasingly prevalent and adapted to the human population as recent outbreaks have spread across the Americas, Caribbean, and Southeast Asia [2–5]. At the peak of the 2016 outbreak, several incidents of imported ZIKV infection were identified in mainland China [6]. In contrast to early and previous epidemics, the recent spread of ZIKV has been associated with severe neurological complications such as Guillain–Barré syndrome in adults and microcephaly in fetuses and newborns [7–10]. Currently, no ZIKV-specific therapeutics or vaccines are available. The high prevalence of the vectors and the continuing evolution of viral species raised serious public health concerns in the near future [11].

The surface envelope glycoprotein (E) of flaviviruses mediates entry and presents a potential target of neutralizing antibodies. Numbers of E-targeting monoclonal antibodies (mAbs) have been identified with potent neutralizing activity and epitope specificity [12–29]. Previously, we isolated and characterized a panel of E-targeting mAbs from plasma and memory B cells from sequential blood samples of a DENV-naïve ZIKV-infected convalescent patient (Pt1) who acquired ZIKV infection in Venezuela during the 2016 outbreak and then returned to China [6, 24]. Among them, ZK2B10 is the most potent in neutralizing ZIKV and have no detectable reactivity with DENV 1 or 2 [24]. ZK2B10 also demonstrated complete prophylactic and impressive therapeutic activities against lethal ZIKV challenge in mouse models of ZIKV infection and microcephaly [30]. Crystal structure and cryo-EM analysis reveal that ZK2B10 recognizes the lateral ridge of DIII and blocks infection at steps between post-attachment and membrane fusion [31]. Since ZK2B10 could serve as a promising candidate for antibody-based interventions, the ontogeny of ZK2B10 could gain insight into the protective antibody response after ZIKV natural infection, as well as inform rational vaccine design. Furthermore, up to date, diverse vaccine candidates are capable of conferring complete protection against ZIKV challenge in mice or nonhuman primates (NHPs) have been evaluated in preclinical and clinical studies [16, 32, 33]. It is therefore imperative to investigate the dynamic and characteristic of antibody repertoire across ZIKV infection longitudinally, which will provide insights into its pathogenesis and the molecular requirement for the development of an effective ZIKV vaccine.

In this study, we applied long-read next-generation sequencing (NGS) and an unbiased repertoire capture method to analyze the B cell repertoire longitudinally of Pt1 from the early acute phase to the late convalescent phase as we previously established [34]. We obtained tens of millions of antibody sequences from a total of seven sequential time points including Day 4, Day 15, Month 2, Month 3, Month 6, Month 10 and Month 12 after the onset of symptoms. We first performed longitudinal analysis of the antibody repertoire, focusing on germline gene usage, CDR3 loop length, and degree of somatic hypermutation (SHM). Our data revealed the antibody repertoire profiles of ZIKV infection with diverse usage of antibody germline gene combined with steady CDR3 loop length, making a clear distinction to chronic HIV infection, which exhibited highly matured repertoire profiles with enrichment of specific germline gene and long HCDR3 loop length [34, 35]. The emerging of germline-like antibodies was observed on Day 15 after symptom onset. We further traced the antibody lineage of ZK2B10 and investigated the maturation pathway. Our results show that ZK2B10 generated in relatively small numbers and grouped with highly germline-like antibodies identified at Day 15 after the onset of symptoms. The somatic variants of ZK2B10 further synthesized for functional characterizations both *in vitro* and *in vivo*. Germline-like heavy chain somatic variants demonstrated strong neutralizing activity and protective potential in lethal ZIKV challenge mouse model. While two substitutions of ZK2B10 λ chain, N31 on LCDR1 and S91 on LCDR3, were identified as the critical residues for ZK2B10 functional maturation. Taken together, our repertoire analyses and lineage tracing elucidated the maturation pathways of potent and protective antibody ZK2B10, and highlighted germline-like antibodies play a noticeable role in protective immunity against ZIKV infection.

## Results

### Dynamic B cell repertoire response throughout ZIKV infection

Next-generation sequencing (NGS) is a powerful tool for probing antibody responses to natural infection and vaccination [36–38]. Extensive studies of broadly neutralizing antibodies (bNAbs) and their lineage development using NGS have revealed the underappreciated complexity and diversity of B cell repertoires in HIV-1-infected individuals during chronic infection [39–42]. Here, we performed a longitudinal NGS analysis of antibody repertoire in Pt1 to delineate the dynamic B cell response to ZIKV infection following the procedure highlighted in Fig 1A. We analyzed seven sequential time points from the acute phase (Day 4 and Day 15 after the onset of symptoms) to the convalescent phase (Month 2, 3, 6, 10 and 12 after the onset of symptoms). We combined 5’-RACE polymerase chain reaction (PCR) and single reverse primers in template preparation to ensure the NGS in an long-read (600 bp) and unbiased manner as previously reported (S1 Fig.) [34, 43–46]. The sequencing yielded a total of 14.2 million heavy chains and 14.1 million light (κ and λ) chains in two separate NGS runs on the Ion S5 GeneStudio platform (S1 Table). The *Antibodyomics* 2.0 pipeline was used to process, annotate, and analyze the NGS data, rendering 1.3 to 2.9 million reads per time point (S1 Table). Of these sequences, 55.3% to 71.2% are high-quality, full-length antibody variable regions which were used for the analyses of B cell repertoire profiles (S1 Table). Furthermore, we traced the lineage of ZK2B10 based on the NGS-derived data and synthesized representative somatic variants for functional characterizations both *in vitro* and *in vivo* (Fig 1A).

**Fig 1.**
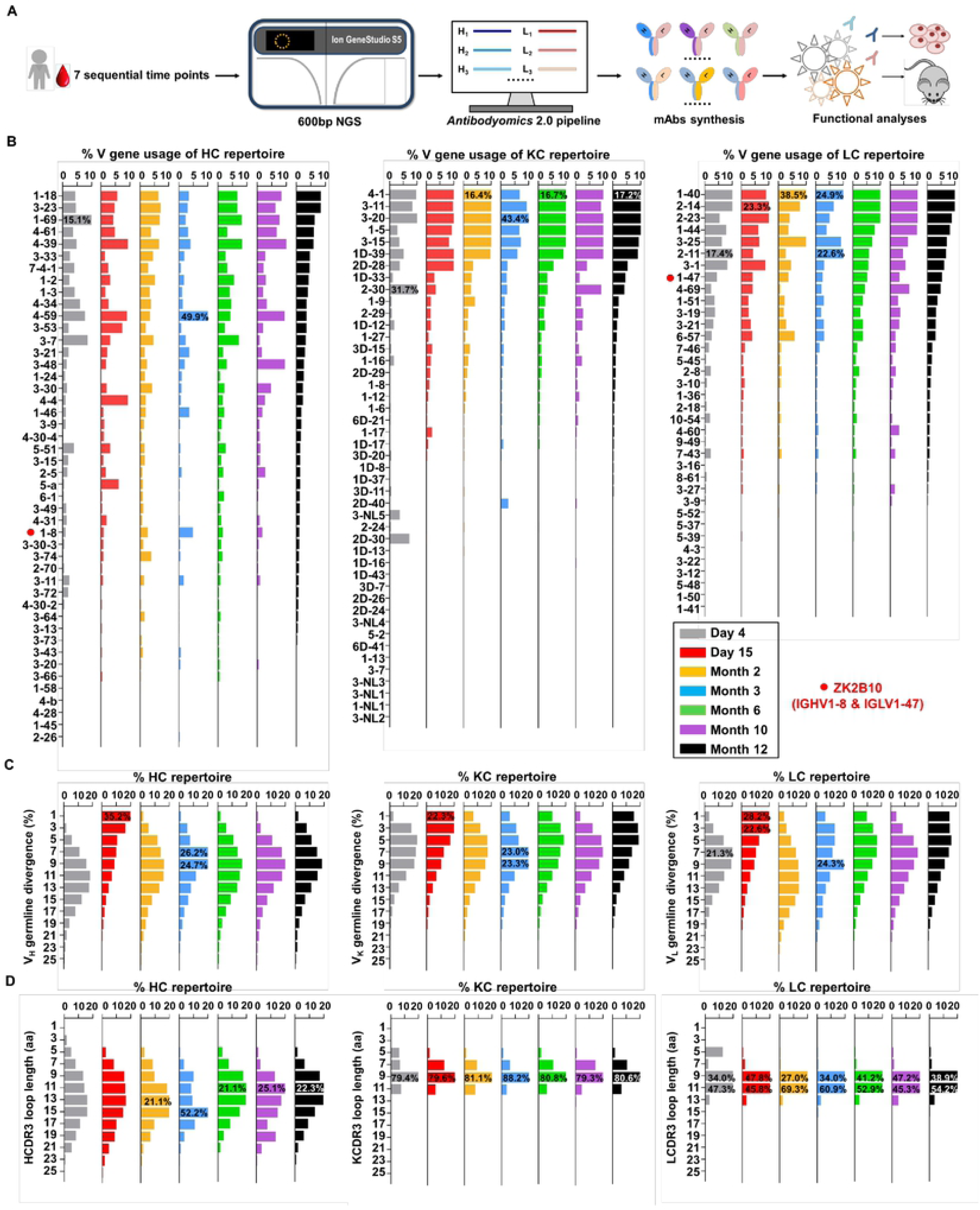
Unbiased antibody repertoire profiles of Pt1 across ZIKV natural infection. **(A)** Schematic view of unbiased antibody repertoire analysis and ZK2B10 lineage tracing. PBMC samples from Pt1 were collected at 7 sequential time points after the onset of symptoms. 5’-RACE PCR was used to prepare antibody chain libraries for long-read (600 bp) next-generation sequencing (NGS) on the Ion GeneStudio S5 platform. The *Antibodyomics* 2.0 pipeline was used to process the NGS data for antibody repertoire profiling, while CDR3-based identification was used for ZK2B10 lineage tracing. Representative somatic variants were synthesized for functional characterizations. **(B-D)** Distributions were plotted for (B) germline V gene usage, (C) germline divergence, and (D) CDR3 loop length for heavy chains (left panel), κ chains (middle panel), and λ chains (right panel). Color coding denotes the 7 sequential time points with Day 4 shown in gray, Day 15 in red, Month 2 in orange, Month 3 in sky blue, Month 6 in green, Month 10 in purple, and Month 12 in black. The germline V genes used by ZK2B10 (IGHV1-8 and IGLV1-47) are labelled in red.

Overall, Pt1 exhibited a diverse and dynamic distribution of germline gene expression (Fig 1B). A few germline genes are dominant in all seven time points such as IGHV1-69, IGKV3-20 and IGLV1-40 with average over 15.21% (Fig 1B, left). In contrast, some specific germline genes were noticed with low frequency, such as IGHV1-8, the V_H_ germline gene of ZK2B10, ranging from 0.98% to 4.80% in seven time points (Fig 1B, left). The V_L_ germline gene of ZK2B10, corresponding to IGLV1-47, ranging from 2.58% to 5.34% (Fig 1B, right). However, the low frequency of IGHV1-8 and IGLV1-47 was unexpected, suggesting that ZK2B10 did not represent a major B cell lineage in the repertoire spanning the acute and convalescent phases of ZIKV infection. In addition, there appeared to be no correlation between the potency of a ZIKV E-targeting mAb and its lineage expansion or prevalence, as indicated by the low frequency of the ZK2B10 germline gene family.

We then determined the degree of somatic hypermutation (SHM) or germline divergence of each time point from early acute phase to late convalescent phase. As shown in Fig 1C, there is a significant increase in the population of germline-like sequences at Day 15 for both heavy and light chains. As a consequence, the average SHM of heavy, κ chain and λ chain repertoire dropped to 6.25%, 5.92% and 5.91% on Day 15, respectively. Of note, the SHM decreased in most V gene families on Day 15 without preference (S2 Fig.). As for the V_H_ germline gene of ZK2B10, IGHV1-8, accounted for only 6.45% SHM on Day 15 and for 7.23% to 13.60% in other time points (S2 Fig., left). The V_L_ germline gene of ZK2B10, corresponding to IGLV1-47, presented 5.72% SHM at Day 15, while 6.70% to 9.04% at other time points studied (S2 Fig., right). These results suggest a drastic shift in repertoire composition likely caused by a rapid plasmablast response during the acute phase of ZIKV infection. The emerging and development of ZK2B10 could represent this kind of antibody response. These patterns corroborate the fact that plasmablasts from ZIKV-infected, flavivirus-naïve individuals exhibited less somatic hypermutation or clonal expansion than those from ZIKV-infected, DENV-immune individuals, which with many derived from common memory B cell clones [19, 47]. Interestingly, the similar pattern has also been reported for an HIV-1 patient undergoing chronic infection in response to a rapidly mutating envelope spike [44].

We next determined changes in the CDR3 loop length. Due to the diversity of the D gene, a rather dispersed distribution in HCDR3 loop length was observed as compared to a steady, canonical CDR3 loop length distribution obtained for κ and λ chains (Fig 1D). The HCDR3 loops were mainly distributed at the range of 9-aa to 15-aa (Fig 1D, left). As for the light chain, 9-aa KCDR3 loops accounted for 79.3% to 88.2% of the κ chain repertoire, while 9-aa to 11-aa LCDR3 loops accounted for 81.3% to 94.9% of the λ chain repertoire (Fig 1D, middle and right). These results revealed a comprehensive view of a human B cell repertoire during ZIKV infection, which revealed the characteristics of acute and transient infection.

### ZK2B10 lineage-specific antibody response during ZIKV infection

To probe the maturation pathway of ZK2B10, we traced the mAb lineage at each time point within the NGS-derived repertoire (Figs 2A and 2B). A CDR3 identity of 95% was used as the cutoff for identifying sequences evolutionarily related to ZK2B10 heavy or λ chain (Figs 2A and 2B, shown as magenta dots on the 2D plots). Unexpectedly, from the library of unbiased amplified germline gene families, we could not find any ZK2B10 heavy chain somatic variants in the repertoire at all seven time points, suggesting that ZK2B10 lineage have an extremely low frequency (Fig 2A, upper panel). To make an in-depth insight into ZK2B10 lineage development, we performed another NGS experiment on four antibody libraries at Day 15, Months 2, 3, and 6, using a degenerate forward primer to target the ZK2B10 heavy chain and its putative germline gene, IGHV1-8 (Fig 2A, lower panel). Gene-specific NGS yielded 1715 ZK2B10-like heavy chains for Day 15 and only two for Month 3 (Fig 2A, lower panel). As for λ chain repertoire, ZK2B10 λ chain somatic variants were detectable on Day 4 but reached the peak on Day 15 with 495 identified variants, and persisted into Month 12 despite a noticeable decline on Month 10 (Fig 2B). Of note, due to the lack of a D gene, light chains do not possess unambiguous sequence signatures for CDR3-based lineage tracing. Nonetheless, our data suggests that ZK2B10 lineage antibodies were induced rapidly and transiently at the end of acute phase during ZIKV infection. Interestingly, the majority of ZK2B10 somatic variants showed a germline divergence of less than 5.0% in both heavy and λ chain repertoires (Fig 2A, low panel and Fig 2B).

**Fig 2.**
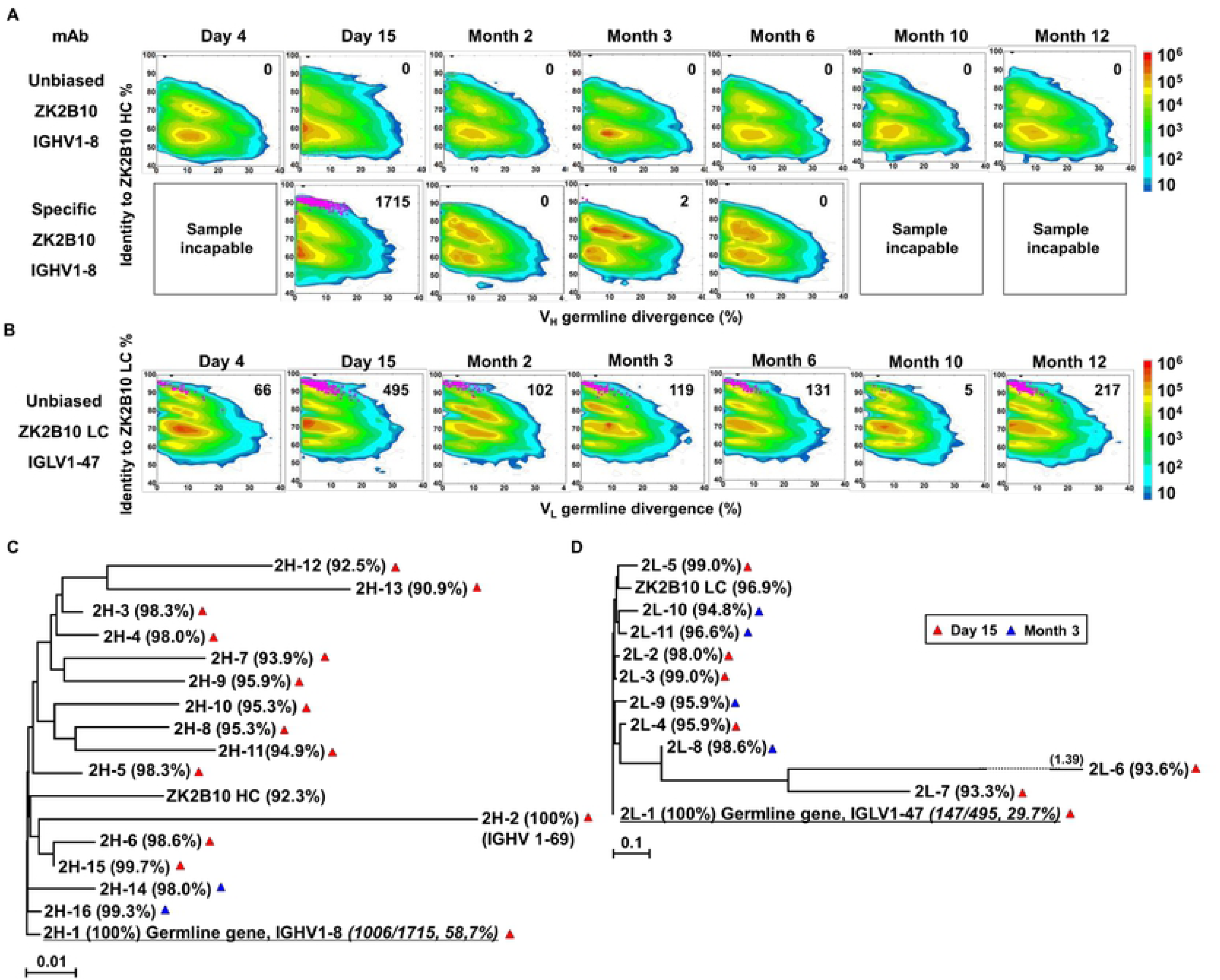
Lineage tracing of ZK2B10 across ZIKV natural infection. **(A)** Lineage tracing of ZK2B10 heavy chain in unbiased amplified library (upper panel) and in ZK2B10-specific amplified library (lower panel). **(B)** Lineage tracing of ZK2B10 λ chain in unbiased amplified library. For each time point, the repertoire is shown as a two-dimensional (2D) plot, with the X-axis indicating the germline divergence of NGS-derived antibody sequences and the Y-axis for their identity with respect to ZK2B10 heavy or λ chain. Color coding indicates the sequence density on the 2D plots ranging from 10^1^ to 10^8^. ZK2B10 heavy or λ chain is shown as block dots on the 2D plots. CDR3-defined somatic variants that evolutionarily related to ZK2B10 heavy or λ chain are shown as magenta dots, with their total number labeled on the 2D plots. HC for heavy chain and LC for λ chain. **(C-D)** Neighbor-joining tree (MEGA6.0) depicting the relationship between germline sequence and the representative somatic variants from ZK2B10 (C) heavy chain and (D) λ chain. Individual variants are named at the branch end point, alongside with their germline identity. Branch lengths are drawn to scale so that the relations between different nucleotide sequences can readily assessed. Color-coding of triangle indicates their emerging time during infection.

To further study the maturation pathway of ZK2B10, we selected representative somatic variants for antibody synthesis and functional characterization. The hierarchical clustering method was used for representative variants selection as previously described [34]. In addition to the dominant sequences, a consensus selection were conducted base on the sequence characteristics to ensure their representativeness [34]. Of these, 16 representative heavy chains were selected with 14 from Day 15 and 2 from Month 3, and 11 representative λ chains with 7 from Day 15 and 4 from Month 3 (Figs 2C and 2D). Surprisingly, among them, 2H-1 and 2L-1 are 100% identical to their putative germline gene, corresponding to IGHV1-8 and IGLV1-47, with sequencing read frequencies as high as 58.7% (1006/1715) and 29.7% (147/495), respectively (Figs 2C and 2D). Furthermore, all these ZK2B10 somatic variants showed a low degree of SHM: the average identity of representative heavy chains with respect to its putative germline gene, IGHV1-8, was 96.81%; as for λ chains, the average germline identity to IGLV1-47 was also as high as 96.79% (Figs 2C and 2D). The variable region sequences and alignments of representative ZK2B10 somatic variants were shown in Fig 3. To summarize, ZK2B10 lineage represents transient plasmablast response with low degree of somatic hypermutation at the end of acute phase of ZIKV infection.

**Fig 3.**
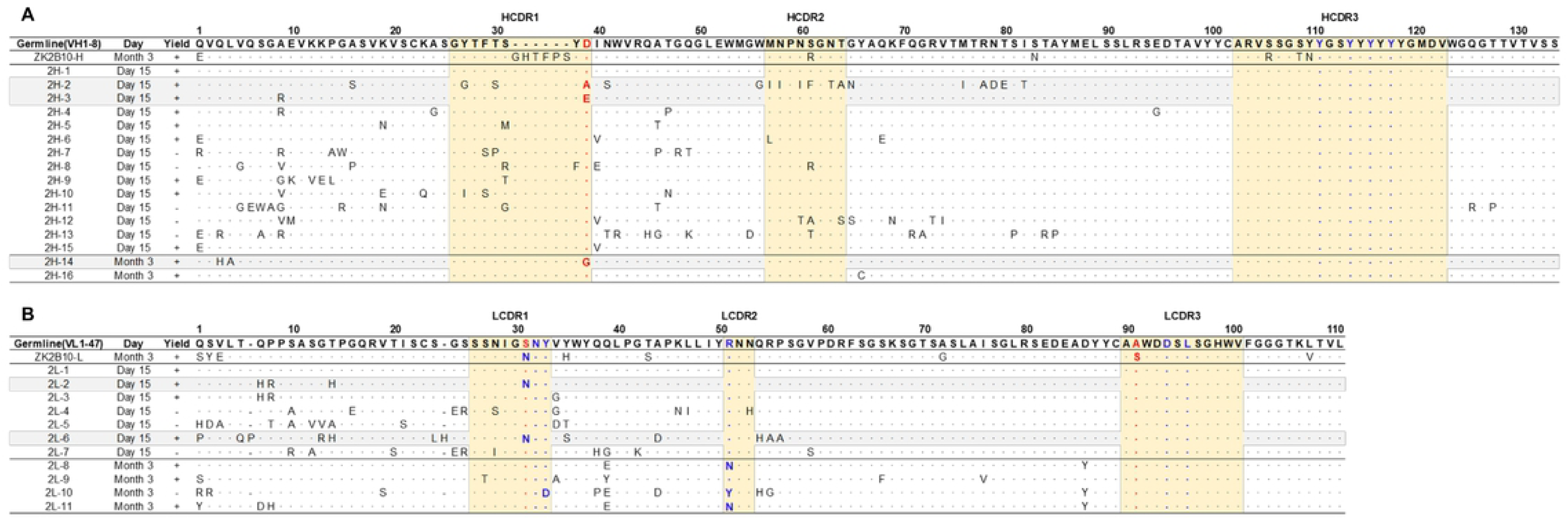
Sequence alignment of representative somatic variants of ZK2B10. **(A-B)** Sequence alignment of representative ZK2B10 (A) heavy chain and (B) λ chain somatic variants with CDR regions highlighted. Residues directly bind to ZIKV E DIII were colored in blue according to the crystal structure of ZK2B10 and ZIKV E DIII complex [31]. The identified potential critical residues for ZK2B10 maturation are marked in red.

### Functional characterization of ZK2B10 somatic variants

The representative ZK2B10 somatic variants were then synthesized and paired with their respective wild-type (WT) partner chains for full-length human IgG1 expression and functional characterization. Of the 16 synthesized ZK2B10 heavy chain somatic variants, 11 (2H-1, −2, −3, −4, −5, −6, −9, −10, −14, −15, and −16) could be expressed when paired with WT ZK2B10 λ chain (Fig 4A). We next measured their binding ability to E glycoprotein and E DIII of ZIKV by ELISA. Among the 11 mAbs, 8 (2H-1, −4, −5, −6, −9, −10, −15, and −16) demonstrated strong binding affinities for E and E DIII at the similar level to ZK2B10, with the half-maximal effective concentrations (EC_50_) ranging from 3.4 to 11.2 ng/ml, while the remaining 3 mAbs (2H-2, −3, and −14) showed weak affinities (Fig 4A). These mAbs were then subjected to a plaque reduction neutralization test against two ZIKV strains, GZ01 (Asian) and MR766 (African), and DENV 2 (Fig 4A). Consistent with their binding abilities, the 8 strong binders (2H-1, −4, −5, −6, −9, −10, −15, and −16) neutralized GZ01 and MR766 potently (Fig 4A). The half-maximal inhibitory concentrations (IC_50_) range from 14.1 to 82.4 ng/ml, which are comparable to WT ZK2B10 and other potent E-targeting mAbs isolated from ZIKV-infected, DENV-naïve human subjects (Fig 4A) [18, 20, 21, 24, 26]. Not surprisingly, 2H-2 failed to show detectable potency (IC_50_ >500 ng/ml to both GZ01 and MR766), 2H-3 showed only modest neutralizing activity (IC_50_= 289.4 ng/ml to GZ01 and 334.1 ng/ml to MR766), and 2H-14 failed to demonstrate high potency against ZIKV as well (IC_50_= 489.1 ng/ml to GZ01 and IC_50_ >500 ng/ml to MR766) (Fig 4A). All these mAbs showed no cross-neutralizing activities with DENV 2 (Fig 4A). Strikingly, with 100% identity to IGHV1-8, 2H-1 showed high affinity for full-length E and E DIII of ZIKV with EC_50_ values of 5.3 ng/ml and 3.4 ng/ml, respectively (Fig 4A). Meanwhile, 2H-1 potently neutralized GZ01 and MR766, with IC_50_ values of 14.1 ng/ml and 19.4 ng/ml, respectively (Fig 4A). Notably, 1006 out of 1715 (58.7%) ZK2B10-like heavy chains from Day 15 were identical to 2H-1, confirming that this germline-like mAb lineage emerged at the peak of plasmablast response. According to the alignment compared to IGHV1-8 germline sequence, the functional loss of 2H-2, −3 and −14 could be explained potentially by the mutation of D39 located toward the end of HCDR1 (Fig 3A). As for 11 synthesized ZK2B10-like λ chains, 7 (2L-1, −2, −3, −6, −8, −9, and −11) were expressible when paired with WT ZK2B10 heavy chain. Surprisingly, 5 (2L-1, −3, −8, −9, and −11) of these 7 λ variants failed to bind ZIKV E or E DIII combined with undetectable neutralizing activities against ZIKV (Fig 4B). Among them, the sequence of 2L-1 is 100% identical to IGLV1-47 and represents a large portion of the Day 15 λ chain population (147 out of 495, 29.7%) (Fig 4B). The reconstituted mAbs containing 2L-2 and 2L-6 demonstrated rather weak binding and neutralizing activities compared to WT ZK2B10 (Fig 4B). Together with the sequence alignments, these patterns presume to that S31N on LCDR1 and in A91S on LCDR3 could be critical for the maturation of ZK2B10 λ chain (Fig 3B).

**Fig 4.**
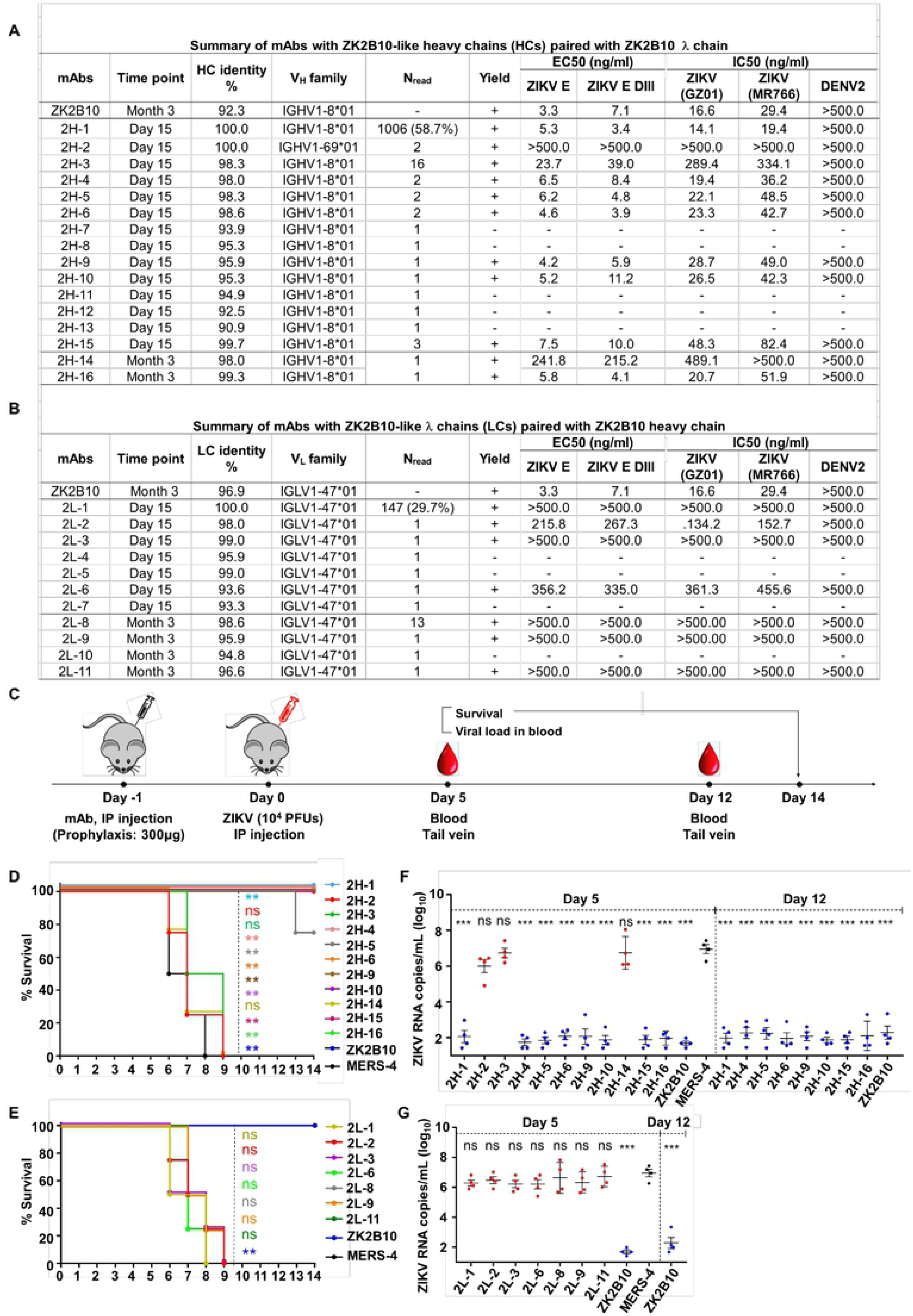
Summary of somatic variants of the ZK2B10 antibody lineage. **(A-B)** ZK2B10 somatic variants are listed with the sampling time point, genetic characterizations, sequencing read frequency (N_read_), expression yield, and functional characterizations. (A) 16 representative ZK2B10 heavy chain variants and (B) 11 representative ZK2B10 λ chain variants identified from the Day 15 and Month 3 antibody repertoires. EC_50_ represents the half-maximal effective concentrations for ELISA binding assays. IC_50_ represents the half-maximal inhibitory concentrations for plaque neutralization assays. **(C-G)** For *in vivo* protection, prophylactic potential against ZIKV infection in AG6 mice was tested. Shown here: (C) timeline for mAb injection, ZIKV inoculation, and blood collection. The prophylactic potential of mAbs was assessed by monitoring survival rates for representative (D) heavy and (E) λ chain somatic variants of ZK2B10 up to 14 days post challenge, and ZIKV RNA copies in blood for (F) heavy and (G) λ chain variants of ZK2B10 on 5 days and 12 days post challenge. Single measurement of ZIKV RNA copies in blood showed distinct results among study groups. Four animals were used in each group. All data are presented as mean ± SEM. *p < 0.05; **p < 0.01; ***p < 0.001; ns, no significant.

To summarize, the results from functional characterizations of ZK2B10 heavy chain somatic variants confirmed the hypothesis that this mAb lineage represents a transient yet effective naïve B cell response to ZIKV infection. The loss of function observed for most λ chain somatic variants, in which V_L_ was reverted to IGLV1-47, suggested that light chain maturation is crucial for the ZK2B10 lineage to acquire its potency and specificity, reminiscent of the case of the HIV-1 bNAb, VRC01 [44].

### Protective potential of ZK2B10 somatic variants in a mouse model

Previously, we have demonstrated ZK2B10 can protect mouse from lethal ZIKV infection and microcephaly [24, 30]. Such animal study will not only confirm the accuracy of our repertoire analyses but also provide clues as to the functional diversity of the ZK2B10 lineage *in vivo*. To this end, we tested the *in vivo* protection against ZIKV lethal infection of representative ZK2B10 somatic variants in AG6 mice (C57BL/6 mice deficient in IFNα, -β, and -γ receptors) following protocol highlighted in Fig 4C [24, 30, 48, 49]. Briefly, we administrated 300μg of each ZK2B10-like mAb, ZK2B10 as positive control, or MERS-4 as negative control to groups of four AG6 mice, from 4 to 6 weeks in age, via the intraperitoneal (i.p.) route (Fig 4C) [50]. On the following day, the animals were challenged with 10^4^ plaque-forming units (PFUs) of ZIKV Asian strain GZ01 via the intraperitoneal (i.p.) route (Fig 4C). Animals were monitored for the survival rate up to 14 days after ZIKV challenge, and for viral RNA level in blood on 5 and 12 days after ZIKV challenge (Fig 4C). As expected, *in vivo* protection of mAbs was correlated with their *in vitro* neutralization, as previously reported [30]. For example, the heavy chain variants with potent neutralizing activities *in vitro* (2H-1, −4, −6, −9, −10, −15 and −16) provided complete protection with a survival rate of 100% up to 14 days after ZIKV challenge (Fig 4D). The RNA load in these groups was suppressed in blood with distinguishable level from the MERS-4 group (Fig 4F). Exceptionally, 2H-2, −3 and −14 failed to offer any protection with a median survival time of 7.25 to 8 days after ZIKV challenge (Fig 4D). The viral RNA levels measured in mice treated by them were 3.72 to 4.45 log_10_ greater on average than the ZK2B10 group on day 5 after ZIKV challenge, respectively (Fig 4F). In contrast, all of the λ chain variants demonstrated an invariant survival rate identical to that of the negative control MERS-4 and failed to suppress viral replication (Figs 4E and 4G). Therefore, *in vivo* evaluation of representative ZK2B10 somatic variants confirmed the differential effect of heavy and λ chains on antibody function, consistent with the *in vitro* characterization by ELISA and neutralization assays.

### Critical residues for ZK2B10 functional maturation

To the further investigation of the maturation pathway of the ZK2B10 lineage, we performed reverse mutagenesis and structural analysis to identify ‘hotspot’ residues. Before that, we aligned the amino acid sequences of representative ZK2B10 heavy and λ chain variants with their putative germline genes, IGHV1-8 and IGLV1-47, respectively (Fig 3). For heavy chain, 2H-2, 2H-3, and 2H-14 lose their potency to ZIKV both *in vitro* and *in vivo.* These 3 heavy chains possess a single substitution mutation at the D39 with respect to their germline gene (Fig 3A). To test that, we conducted reverse mutagenesis on 2H-2 (A39D), 2H-3 (E39D), and 2H-14 (G39D) and characterized the function of these mutants by ELISA and neutralization assays. As shown in Fig 5A, 2H-3 (E39D) and 2H-14 (G39D) mutants regained their ZIKV E-binding and neutralizing activities, approaching the level of WT ZK2B10. Due to the use of a different V_H_ germline gene, IGHV1-69, the 2H-2 (A39D) mutant was ineffective (Fig 5A). This result suggested that the ZK2B10 lineage has a restricted V_H_ gene usage to achieve high affinity and potency against ZIKV. Based on the crystal structure of ZK2B10 in complex with ZIKV E DIII, 4 residues within HCDR3 loop (Y111, Y114, Y116 and Y118) were directly involved in the contact interface [31]. Although D39 is on HCDR1 loop, it forms hydrogen bonds with Y115 and Y117 on the opposite side of the HCDR3 loop, thus stabilizing the HCDR3 conformation (Fig 5C). These results provide further evidence that the ZK2B10 lineage was indeed generated during the naïve B cell response to acute ZIKV infection, with critical residues encoded by the germline gene. For λ chain variants, two critical mutations were identified that potentially contribute to the maturation of ZK2B10 lineage (Fig 3B). One such mutation, located on the LCDR1 loop, is N31, which is shared by the weak-functional variants 2L-2 and 2L-6, as well as WT ZK2B10 λ chain (Fig 3B). The other is at position 91 on LCDR3 loop, which is S91 in WT ZK2B10 λ chain but predominantly A91 in all these weak- or non-functional λ chain variants (Fig 3B). Thus, N31N and S91 could be the critical mutations for ZK2B10 λ chain maturation. We first examined the effect of these two mutations individually by performing site-directed mutagenesis on 2L-1, which is 100% identical to germline gene IGLV1-47. Neither S31N nor A91S could effectively render the germline antibody functional (Fig 5B). We then introduced a double mutation (S31N+A91S) into 2L-1, which, as expected, bound to ZIKV E and E DIII with high affinity and potently neutralized ZIKV at the same level of WT ZK2B10 (Fig 5B). As shown by the crystal structure, 5 residues within ZK2B10 λ chain (N31, N32, Y33, R51, D94 and L96) directly involved in the in the contact interface [31]. For the two identified critical residues, N31 directly interacts with T309 on ZIKV E DIII, while S91 forms a hydrogen bond with N32, which interacts with T335 on ZIKV E DIII (Fig 5D). In brief, our combined analyses of NGS data, antibody functions, and complex structure confirm that residues N31 and S91 within λ chain are essential to the function of ZK2B10, thus posing a major barrier to the functional maturation of this ZIKV E DIII-directed antibody lineage.

**Fig 5.**
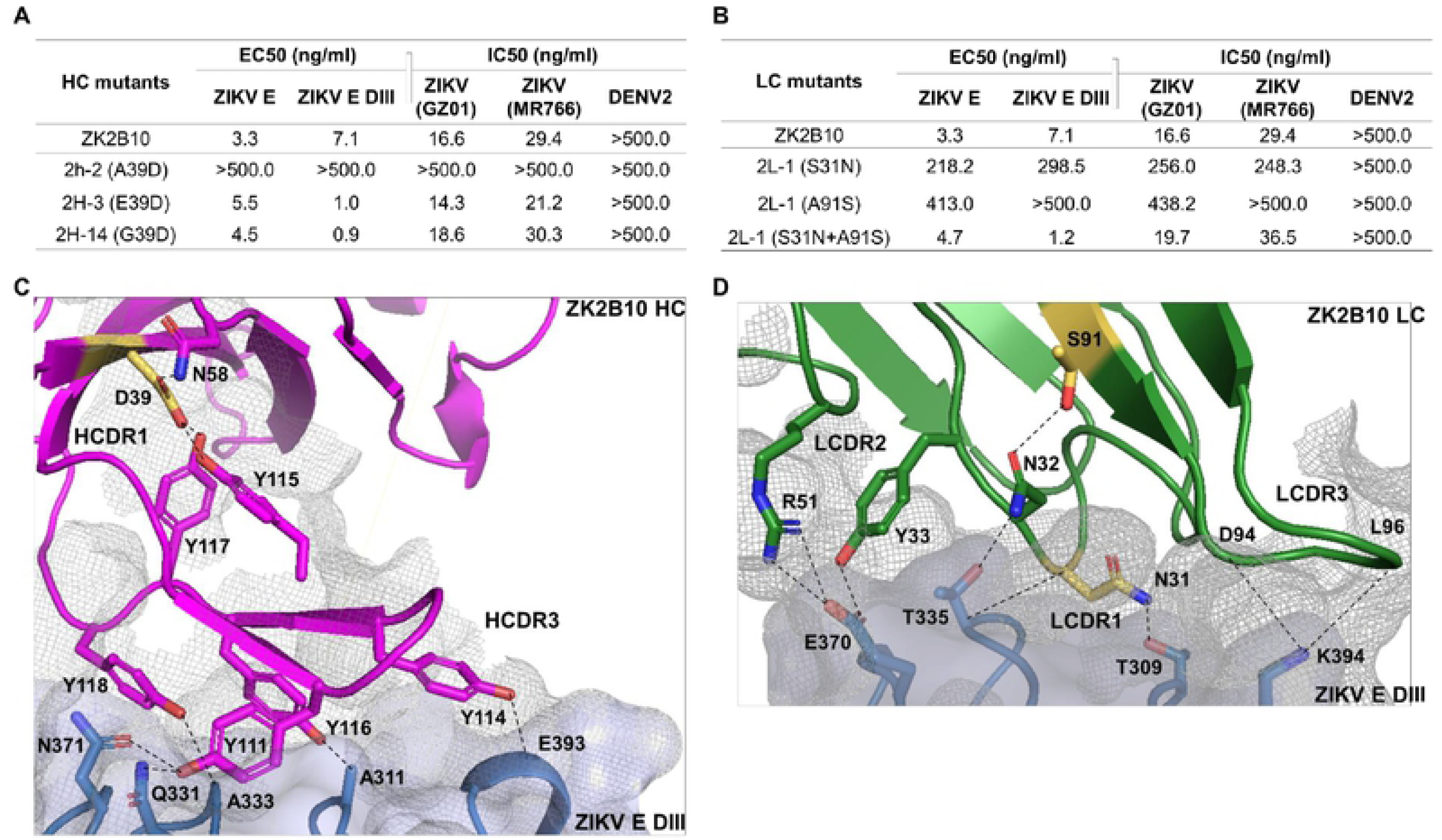
Mutagenesis and structural analyses of critical residues for ZK2B10 maturation. **(A)** Validation of ZK2B10 heavy chain critical residues by reverse mutagenesis. **(B)** Validation of ZK2B10 λ chain critical residues by mutagenesis. **(C)** The contact interface of ZK2B10 heavy chain (HC) with ZIKV E DIII. In the ribbon diagram, the ZK2B10 HC is shown in magenta and ZIKV E DIII in cornflower blue. Residues directly involved in the contact interface with E DIII (Y111, Y114, Y116 and Y118) are shown with side chain. The identified critical residues, D39 on HCDR1, is highlighted in yellow. **(D)** Contact interface of ZK2B10 λ chain (LC) with ZIKV E DIII. ZK2B10 LC is shown in forest green and ZIKV E DIII in cornflower blue. Residues directly involved in the contact interface with E DIII (N31, N32, Y33, R51, D94 and L96) are shown with side chain. The identified critical residues, N31 on LCDR1 and S91 on LCDR3, are highlighted in yellow.

## Discussion

In this study, we delineated the human B cell repertoire profiles across ZIKV natural infection by accurate NGS approach. The comprehensive analyses showed antibody repertoire profiles with diverse germline usage, lower IgG somatic hypermutation rate and steady CDR3 loop length. The tracking of ZK2B10 revealed the dynamic of an effective germline-coded antibody lineage, which emerged preluding the convalescent phase of ZIKV infection. Germline-like somatic variants derived from ZK2B10 lineage neutralized ZIKV potently and protected mice from lethal challenge, while demonstrated no cross-reactivity with DENV2. The in-depth analyses showed that two site-mutagenesis of IGLV1-47 germline-coded λ chain, N31 and S91, are essential to the functional maturation of IGHV1-8/IGLV1-47 antibody lineage.

Two unique aspects of our study are worth highlighting here. One is based on the effective germline-coded antibody response represented by ZK2B10 lineage. We report here that there was a significant increase in germline-like antibodies at Day 15 after the onset of symptoms. This drastic shift in repertoire composition was likely a result of rapid plasmablast response toward the end of the acute phase of ZIKV infection. Interestingly, this shift was coincided with the emergence of the ZK2B10 lineage, which provides a perfect introduction to understand the germline-coded antibody response during ZIKV natural infection. Similar patterns have been reported in separate antibody studies. For monoclonal antibodies, germline-like human mAbs, m301 and m302, were reported that target ZIKV E DIII cryptic epitopes (C-C’ loop) and neutralize ZIKV potently both *in vitro* and *in vivo* [22]. Another human mAb, P1F12, originates from germline gene IGHV3-7 with an identity of 100% and neutralizes ZIKV potently as well [51]. For the overall B cell response, plasmablast-derived antibodies from ZIKV-infected, DENV-naïve donor showed low levels of SHM, supporting the mechanism of naïve B cell activation [19]. Lower IgG somatic hypermutation rates also reported during acute DENV infection, which is consistent with an innate-like antiviral recognition mediated by B cells using defined germline-coded B cell receptors [52]. Therefore, the inducing of germline-coded neutralizing antibodies occupies critical positon in the B cell response during acute flavivirus infection. Diverse mechanisms of antibody lineage development were found in chronic infection, especially in HIV studies. The initiation and early development of MPER-directed HIV antibody lineage have been reported to achieve high neutralizing breadth with low mutation from germline [53]. In a stark contrast, HIV-1 bNAbs VRC01 and PGT121, which target CD4 binding-site and V3 region respectively, require extensive mutation to achieve neutralizing breadth and potency and possess long HCDR3 loops to penetrate the glycan shield of the HIV-1 envelope spike [34, 44]. In summary, our longitudinal analysis of antibody repertoire and lineage development provides critical insights into the pathogenesis of ZIKV infection.

The other unique aspect of our study was the assist in the rational design of a safe and effective vaccine. As we previously reported that ZK2B10, as well as other E DIII-specific mAbs, are ZIKV-specific, potent neutralizing and protective in mice from a lethal ZIKV challenge [18, 23, 24, 30]. The identified epitope reveal that ZK2B10 bind to residues within the lateral ridge of DIII and blocks infection at a post-attachment step as other E DIII-specific potent neutralizing mAbs [31, 47]. Beyond that, DIII-specific antibodies give essential contribution to controlling ZIKV as they correlated positively with high neutralization titers and the depletion of them results in reduced neutralizing activity in ZIKV-infected patient serum [24, 47]. We also report here that only two site-mutagenesis of IGLV1-47 germline λ chain, N31 and S91, are sufficient for IGHV1-8/IGLV1-47 germline antibodies to achieve potent ZIKV neutralization. This barrier could overcome readily by an active B cell repertoire. The low degree of SHM observed for the ZK2B10 lineage suggests that elicitation of naïve protective B cell response against ZIKV may be achievable with a standard vaccination regimen. Furthermore, E DIII-based vaccine has been reported to avert lethal West Nile virus (WNV) infection without enhancing ZIKV or DENV infectivity [54]. However, the low frequency and transient expansion of ZK2B10-like antibodies in the Pt1 repertoire suggest that overcoming the suboptimal immunogenicity of ZIKV E DIII, an elongated immunoglobulin-like domain, may prove to be a challenge for ZIKV vaccine development [55, 56].

## Materials and methods

### Donor and PBMCs samples

The blood samples were donated by a 28-year-old Chinese ZIKV convalescent male patient (Pt1) who traveled from Venezuela to the southern metropolitan city Guangzhou, China, in February, 2016 [6]. During his hospitalization and follow-up visits, a total of 7 sequential blood samples were collected on Day 4, Day 15, Month 2, Month 3, Month 6, Month 10 and Month 12 after the onset of the symptoms. Samples were separated into plasma and peripheral blood mononuclear cells (PBMCs) by centrifugation through a Ficoll-Hypaque gradient (GE Healthcare). PBMCs were cryopreserved in freezing media and stored in liquid nitrogen until further analysis by antibody repertoire sequencing.

### Sample preparation using 5**’**-RACE PCR

An improved version of the 5’-RACE PCR protocol for sample preparation is reported in a recent study [34, 44]. Here, total RNA was extracted from 1∼5 million PBMCs into 30ml of water with RNeasy Mini Kits (Qiagen, Valencia, CA). For unbiased repertoire analysis, 5’-RACE was performed with SMARTer RACE cDNA Amplification Kit (Clontech, Mountain View, CA). For ZK2B10 gene-specific lineage analysis, reverse transcription (RT) was performed with SuperScript III (Life Technologies) and oligo (dT). In both cases, the cDNA was purified and eluted in 20ul of elution buffer (NucleoSpin PCR Clean-up Kit, Clontech). The immunoglobulin PCRs were set up with Platinum Taq High-Fidelity DNA Polymerase (Life Technologies, Carlsbad, CA) in a total volume of 50 µl, with 5 μl of cDNA as template, 1 μl of 5’-RACE primer or gene-specific forward primers, and 1 μl of 10 µM reverse primer. To facilitate deep sequencing on the Ion GeneStudio S5 system, the forward primers (both 5’-RACE and gene-specific) contained a P1 adaptor, while the reverse primer contained an A adaptor and an Ion Xpress^TM^ barcode (Life Technologies) to differentiate the libraries from various time points. A total of 25 cycles of PCRs were performed and the PCR products (∼600 bp for 5’-RACR PCR or ∼500 bp for gene-specific PCR) were gel purified (Qiagen, Valencia, CA). A degenerate primer (SAGGTGCAGCTGGTGCAGTCTGG) was used as the forward gene-specific primer to cover potential variations at the 5’-end of ZK2B10 transcripts.

### Next-generation sequencing (NGS) and *Antibodyomics* analysis

Antibody NGS has been adapted to the Ion GeneStudio S5 system [57]. Briefly, the antibody heavy and light (κ and λ) chain libraries were quantitated using Qubit® 2.0 Fluorometer with Qubit® dsDNA HS Assay Kits. Equal amounts of the heavy chain libraries from various time points were mixed and loaded onto an Ion 530 chip to increase the sequencing depth and to eliminate run-to-run variation. The κ and λ chain libraries at each time point were mixed at a ratio of 1:1 prior to library pooling and chip loading. Template preparation and (Ion 530) chip loading was performed on the Ion Chef system using Ion 530 Ext Kits, followed by S5 sequencing with the default settings. Raw data was processed without 3’-end trimming in base calling to extend the read length. The human *Antibodyomics* pipeline version 1.0 [34, 39, 44] has been modified to improve data accuracy and computational efficiency [57]. This new *Antibodyomics* pipeline was used to process and annotate Pt1 antibody NGS data for repertoire profiling and lineage tracing. The distributions of germline genes, germline divergence or degree of somatic hypermutation (SHM), and CDR3 loop length derived from antibody NGS data as general repertoire profiles. The two-dimensional (2D) divergence/identity plots were constructed to visualize ZIKV-specific antibody lineages in the context of Pt1 antibody repertoire. A CDR3 identity of 95% was used as the cutoff for identifying sequences evolutionarily related to a reference antibody (shown as magenta dots on the 2D plots). The hierarchical clustering method was used to divide CDR3-defined somatic variants into groups based on an overall identity cutoff of 98% as previously described [34]. In addition to the dominant sequences, a consensus or a manually selected sequence was used as the group representative for antibody synthesis and functional characterization. ZK2B10 were initially isolated from PBMCs of Pt1 as we previously reported [24].

### Human monoclonal antibody (mAb) clones construction, expression, and purification

All of the synthetic variable region genes of antibody heavy chain (V_H_) and light chain (V_K/L_) were analyzed using the IMGT/V-Quest server (http://www.imgt.org/IMGTindex/V-QUEST.php). They were cloned into the backbone of antibody expression vectors containing the constant regions of human IgG1 as previously described [50]. To produce full-length human mAbs, the recombinant clone was paired with the complementary chain of wild-type (WT) ZK2B10. The heavy and light chain expression plasmids were transiently co-transfected into HEK 293T cells for the production of full-length human IgGs, which were purified from the supernatant by affinity chromatography using protein A agarose (Thermo Scientific). The IgG concentration was determined using the BCA Protein Assay Kit (Thermo Scientific). We included previously reported MERS-CoV-specific mAb MERS-4 [50] for comparative analysis.

### ZIKV E and ZIKV E DIII protein and ELISA analyses

The genes of either E protein or E DIII protein (residues 301-403) of ZIKV (GZ01, KU820898) without tag were cloned into pET28a vectors (Novagen) and expressed by IPTG-induction in BL21 (RIL) bacterial cells. The isolated inclusion bodies were solubilized and re-folded as reported [55]. In ELISA binding assays, the E proteins and E DIII proteins were captured separately onto ELISA plates overnight at 4 °C. Each tested mAb was serially diluted and applied to the ZIKV E and E DIII protein-captured ELISA plates. Binding activities were detected using anti–human IgG labeled with HRP and TMB substrate.

### Antibody neutralization assays

All ZIKV GZ01 (KU820898), ZIKV MR766 (AY632535) and DENV2 43 (AF204178) viruses were grown in C6/36 Aedes Albopictus cells and titrated on Vero cells before use. For neutralization assay, serial dilutions of mAbs were mixed with virus at 4 °C for 1 hour before being applied to Vero cells in the 6-well culture plates. After 1–2 hour of infection, the antibody-virus mixture was aspirated and Vero cells were washed with PBS and overlaid with DMEM containing 2% heat-inactivated FBS and 1% SeaPlaque Agarose (Lonza, 50501). After 4–6 days, plaques were stained by 1% crystal violet and counted manually.

### Antibody prophylactic potential analyses in AG6 mice

C57BL/6 mice deficient in interferon (IFN) α, -β, and -γ receptors (AG6 mice) were kindly provided by the Institute Pasteur of Shanghai, Chinese Academy of Sciences (IPS). The mice were bred and maintained in a pathogen-free animal facility. Groups of 4 sex-matched, 4-to 6-week-old AG6 mice were used for the animal studies. In prophylaxis assays, 300μg of each tested mAb or isotype control (MERS-4) was administered via the i.p. route. The following day, the animals were challenged with 10^4^ PFUs of ZIKV (GZ01 strain) via i.p. injection. Survival were monitored for up to 14 days post challenge. On days 5 and 12 after challenge, whole blood was collected from each animal for ZIKV viral load measurement.

### Quantitative measurement of viral loads by TaqMan qPCR

Whole blood (10 μL) was collected in an RNase free Eppendorf tube containing lysis buffer (QIAGEN) and stored at −80°C until use. Total RNA was extracted using RNeasy Mini Kits (74106, QIAGEN) and reverse-transcribed into cDNA using iScript cDNA Synthesis Kits (170-8890, Bio-Rad). Viral RNA copies were quantified through TaqMan qPCR amplification of ZIKV (GZ01) envelope gene. Measurements were expressed as log_10_ viral RNA copies per millimeter calculated against a standard curve. Sequences for primers and probes were as follows: ZIKV-F CCGCTGCCCAACACAAG, ZIKV-R CCACTAACGTTCTTTTGCAGACAT, ZIKV-probe AGCCTACCTTGACAAGCARTCAGACACTCAA (5’FAM, 3’TAMRA)

### Multiple sequence alignment and structural analysis

Multiple sequence alignment (MSA) was calculated using BioEdit ClustalW. The crystal structure of ZIKV DIII-ZK2B10 Fab complex has been determined and analyzed here to identify the ‘hotspot’ residues critical to ZK2B10 lineage development [31]. For the intermolecular interactions shown in Fig 5, 4 Å was used as the maximal cut-off distance for hydrogen bonds. Illustrations of structural models were prepared using PyMOL Molecular Graphics System 1.5.0.4.

### Statistical methods

All data were analyzed using Prism6 software (GraphPad). The half-maximal effective concentrations (EC_50_) were calculated using the dose-response stimulation model. The IC_50_ value for each mAb was calculated using the dose-response inhibition model. For experiments involving AG6 mice, 4 animals were included in each assessment group to ensure equal representation and consistency of the data obtained. Statistical evaluation was performed using Student’s unpaired t test. Data were presented as mean ± SEM. *p < 0.05; **p < 0.01; and ***p < 0.001.

### Ethics statement

The human study was approved by the Ethical Committee of the Guangzhou Eighth People’s Hospital, Guangzhou Medical University. The research was conducted in strict accordance with the Chinese government rules and regulations for the protection of human subjects. The study subjects provided the written informed consents for research use of their blood samples. All procedures with animals were undertaken according to Experimental Animal Welfare and Ethics Committee of Tsinghua University. All experiments were performed under the guidelines of the Experimental Animal Welfare and Ethics Committee of Tsinghua University (16-ZLQ9).

## Acknowledgements

We are grateful to the ZIKV convalescent patient for donating his blood samples from which mAbs were isolated and Drs. Jiang Wang, Wenxin Hong, and Lingzhai Zhao for providing the patient with treatment and care. We thank Drs. Cheng-Feng Qin and Gong Cheng for providing the Zika virus isolate GZ01 and MR766. We are also grateful for Institute Pasteur of Shanghai, Chinese Academy of Sciences for kindly providing AG6 mice.

## Author contributions

Project design by F.G., X.L., L.Z., J.Z, L.Y.; sample preparation by F.G. and L.Z.; library preparation and NGS by L.H.; data processing and annotation by X.L. and J.Z.; antibody lineage tracing by X.L. and J.Z.; antibody sequence selection by F.G., X.L., and J.Z.; antibody synthesis by F.G. and L.Z.; antigen binding and neutralization assays by F.G. and L.Z.; ZIKV challenge and protection in mice by F.G. and L.Z.; manuscript written by F.G., L.Z., J.Z., and L.Y.

---

**S1 Fig. Strategy for Next Generation Sequencing and CDR3-based lineage tracing.** To facilitate NGS, 5’ RACE PCR was used for unbiased antibody repertoire analysis, while ZK2B10 heavy chain specific amplification primer was for ZK2B10 HC lineage tracing. The forward primers contained a trP/P1 adaptor, while reverse primer contained an A adaptor and a N_10_ barcode to differentiate the libraries of various time points. To improve data accuracy, the raw sequences of antibody heavy chain or light (κ/λ) chain variable regions were processed separately by *Antibodyomics* pipeline. After CDR3-based identification, somatic variants with 95% or greater CDR3 identity on nucleotide level were defined for lineage tracings. Representative somatic variants were synthesized for full-length human IgG1 production followed by further *in vitro* and *in vivo* functional assays.

**S2 Fig. Distribution of germline divergence in each V gene germline of Pt1 across ZIKV natural infection.** Distributions were plotted for the germline divergence in each V gene germline of heavy chains (left panel), κ chains (middle panel) and λ chains (right panel). Color coding denotes the 7 sequential time points with Day 4 shown in gray, Day 15 in red, Month 2 in orange, Month 3 in sky blue, Month 6 in green, Month 10 in purple, and Month 12 in black.

**S1 Table. Next-generation sequencing (NGS) of antibody repertoires of a ZIKV-infected Chinese patient.**

